# The genetic and environmental etiology of blood-based biomarkers related to risk of Alzheimer’s Disease in a population-based sample of early old-age men

**DOI:** 10.1101/2022.08.09.503234

**Authors:** Nathan A. Gillespie, Robert A. Rissman, Jeremy A. Elman, Ruth E. McKenzie, Xin M. Tu, Hong Xian, Chandra A. Reynolds, Matthew S. Panizzon, Michael J. Lyons, Graham M.L. Eglit, Michael C. Neale, Carol Franz, William S. Kremen

## Abstract

The amyloid-tau-neurodegeneration (ATN) framework has led to an increased focus on Alzheimer’s disease (AD) biomarkers. The cost and invasiveness of obtaining biomarkers via cerebrospinal fluid has motivated efforts to develop sensitive blood-based biomarkers. Although AD is highly heritable, the biometric genetic and environmental etiology of blood-based biomarkers has never been explored. We therefore, analyzed plasma beta-amyloid (Aβ40, Aβ42, Aβ42/40), total tautau (t-tautau), and neurofilament light (NFL) biomarkers in a sample of 1,050 men aged 60 to 73 years (m=68.2, SD=2.5) from the Vietnam Era Twin Study of Aging (VETSA). Unlike Aβ and tautau, NFL does not define AD; however, as a biomarker of neurodegeneration it serves as the N component in the ATN framework. Univariate estimates suggest that familial aggregation in Aβ42, Aβ42/40, t-tau, and NFL is entirely explained by additive genetic influences accounting for 40%-58% of the total variance. All remaining variance is associated with unshared or unique environmental influences. For Aβ40, a additive genetic (31%), shared environmental (44%), and unshared environmental (25%) influences contribute to the total variance. In the more powerful multivariate analysis of Aβ42, Aβ40, t-tau, and NFL, heritability estimates range from 32% to 58%. Aβ40 and Aβ42 are statistically genetically identical (r_g_ = 1.00, 95%CI = 0.92,1.00) and are also moderately environmentally correlated (r_e_ = 0.66, 95%CI = 0.59, 0.73). All other genetic and environmental associations were non-significant or small. Our results suggest that plasma biomarkers are heritable and that Aβ40 and Aβ42 share the same genetic influences, whereas the genetic influences on plasma t-tau and NFL are mostly unique and uncorrelated with plasma Aβ in early old-age men.

## Introduction

Alzheimer’s disease (AD) is the most costly disease in the U.S. ^[80]^, particularly in terms of the years of life lost and the years lived with disability ^[29]^. With the failure of recent drug trials and recognition that the disease process in AD begins decades before dementia onset, there is now widespread consensus that early identification is key to preventing or slowing disease progression ^[16, 19, 22–24]^. The protracted prodromal period in AD also calls for a focus on earlier identification with regard to cognitive decline, mild cognitive impairment (MCI), and preclinical signs of AD ^[19, 23, 24, 32]^. Arguably, addressing early risk factors could be a step toward the “ounce of prevention” that would be “worth a pound of cure.” Estimates are that a 5-year delay of the dementia phase of AD would reduce the number of cases by half ^[24]^. Moreover, the public health impact of such delays will only grow in the next decade with the increasing number of 65 to 75-year-olds ^[25]^.

Biomarkers are central to the definition of AD ^[59]^ and given the increasing emphasis placed on detecting earlier those individuals who are at risk, plasma-based biomarkers have come under increasing attention. The advantages of plasma biomarkers include accessibility and affordability. Unfortunately, only a fraction of brain protein enters the bloodstream making biomarkers difficult to measure. Moreover, dilution, degradation, or metabolism introduces variance unrelated to AD-related brain changes that is difficult to control. These factors might limit the predictive validity of biomarkers earlier in life ^[37, 51, 66]^. Fortunately, innovative developments relying on ultrasensitive immunoassays and novel mass spectrometry techniques have begun to show promise in terms of leveraging plasma biomarkers to measure beta-amyloid (Aβ40, Aβ42, and the Aβ42/40 ratio) and tau, the two hallmark pathologies of AD, and neurodegeneration (tau and neurofilament light proteins) ^[37, 51, 66]^.

Amyloidosis may offer predictive validity in terms of AD pathophysiology. For example, low plasma Aβ levels may identify adults with MCI and AD ^[13, 35, 38, 39]^ including cases 8 to 15 years before dementia onset ^[15, 18, 27, 28, 36, 38]^. Nakamura et al. ^[60]^ have reported robust correlations between plasma biomarkers and areas of high Aβ deposition in the brain. Levels of plasma Aβ40, Aβ42, and the Aβ42/40 ratio are all significantly correlated with cerebrospinal fluid (CSF) Aβ when analyses are based on AD cases and controls, subjects with MCI and subjective cognitive decline (SCD) ^[43]^. Levels of Aβ42 and the Aβ42/40 ratio from plasma are also significantly correlated with amyloid positron emission tomography (PET) standardized uptake value ratio when examined across all cases (SCD, MCI, and AD) ^[43]^. We note, however, that findings regarding plasma Aβ are equivocal; several reports have shown either no association between Aβ biomarkers and AD pathophysiology or mixed findings depending on the particular Aβ marker ^[44, 53, 63, 69]^. For example, Janelidze et al. ^[43]^ reported that *APOE*-ε4 carriers show significantly lower levels of Aβ42 (p < 0.001), Aβ40 (p = 0.009) and a lower Aβ42/40 ratio in plasma compared to non-carriers. However, when analyzed within individual diagnostic groups, plasma Aβ42 was decreased in *APOE*-ε4 carriers in controls and individuals with SCD, but not among those with either MCI or AD. Plasma biomarkers might prove to be a useful tool for monitoring synaptic degeneration and AD pathophysiology ^[66]^ or for screening individuals in the prodromal stages of AD ^[43, 74, 75]^.

Tau proteins are a group of six highly soluble protein isoforms produced by alternative splicing from the *MAPT* gene ^[3, 4]^. In AD patients, tau loses the ability to bind to microtubules and therefore its normal role of keeping the cytoskeleton well-organized is no longer effective ^[26]^ and is abnormally hyperphosphorylated but without ubiquitin reactivity ^[49]^. Associations between plasma tau and AD pathophysiology have, however, yielded equivocal results. For example, individual studies have reported either no association between plasma and CSF t-tau ^[63]^ or elevated but nonsignificant associations, e.g., between plasma and PET t-tau in AD dementia patients ^[59]^. Fiandaca et al. ^[40]^ found that combining plasma phosphorylated tau (p-tau) and Aβ42 yielded a 96% sensitivity for differentiating AD and MCI groups from cognitively normal older adults. Olsson et al.’s ^[44]^ meta-analysis of 231 reports found that plasma t-tau was significantly associated with AD. T-tau has been used as a marker of general neurodegeneration whereas p-tau is considered to reflect the formation of neurofibrillary tangles^[77].^

Plasma and CSF NFL concentrations have been shown to be highly correlated (r = 0.59 to 0.89) ^[41, 48]^. Plasma NFL is significantly increased in individuals with MCI and patients with AD dementia when compared to controls ^[48]^. Plasma NFL significantly correlates with functional scores in patients with behavioral variant frontotemporal dementia and the functional performance of patients with AD and MCI ^[62]^. Baseline levels of NFL are also higher in patients with MCI and AD dementia compared to controls ^[70]^.

Mattsson et al. ^[70]^ have argued that plasma NFL is a noninvasive biomarker linked to neurodegeneration in patients with AD. Likewise, Preische et al. ^[76]^ have argued that because changes in plasma NFL predict disease progression and brain neurodegeneration during the early pre-symptomatic stages of familial AD ^[76]^, NFL’s utility as an AD biomarker is supported ^[76]^. In contrast, Blennow and Zetterberg’s ^[66]^ review argued that plasma NFL is not a feature specific to AD, but is found in many neurodegenerative disorders, and thus should only be employed as a screening tool for subjects with cognitive disturbance to rule out neurodegeneration. In any case, as a biomarker of neurodegeneration, NFL can be a useful indicator of the N component of the ATN framework.

While research using improved mass spectrometry techniques to test the validity of plasma biomarkers continues ^[66]^, remarkably little is known about the genetic and environmental etiology of these biomarkers or their sources of covariation. We are aware of small-sampled genome-wide association studies examining CSF Aβ or tau-protein species ^[17, 46, 54, 72, 81]^, plasma Aβ ^[34]^, plasma NFL ^[58]^, plasma tau ^[45]^ and one whole-exome sequence-based association study ^[50]^ examining Aβ42/40. However, given the small sample sizes none of these molecular reports reported either SNP heritability or genetic correlations between biomarkers. We are unaware of any twin studies that have examined the etiology of either CSF or plasma-based AD biomarkers. Another limitation is that biomarker studies typically rely on elderly adult samples, or clinically ascertained subjects (e.g., from memory clinics), individuals with high SES, or cross-sectional comparisons between AD cases and controls that can be confounded by genetic and environmental differences.

We addressed these limitations by exploring the genetic etiology of AD plasma biomarkers in a large community-dwelling sample of early old-age male twins from whom we obtained blood-plasma. Our specific aims included estimating i) the standardized contribution of genetic and environmental influences in Aβ40, Aβ42, Aβ42/40, t-tau and NFL, and ii), the genetic and environmental correlations between them. To the extent that plasma-based biomarkers are unreliable, the expectation is that individual differences will be largely explained by random environmental variance that includes measurement error. However, if plasma-based biomarkers are indeed capturing reliable variation, there may be significant familial aggregation in the form of either genetic or shared environmental influences.

## Materials and methods

### Subjects

The Vietnam Era Twin Study of Aging (VETSA) is a longitudinal study of cognitive and brain aging and risk for Alzheimer’s disease in a national US sample of community-dwelling men ^[31]^. The present study comprised those who participated in the third assessment wave when plasma biomarkers were examined. Briefly, Wave 1 took place between 2001 and 2007 ^[12]^ (mean age=55.9, SD=2.4, range=51.1 to 60.7). Wave 2 occurred approximately 5.5 years later (mean age=61.7, SD=2.5, range=56.0 to 67.0). Wave 3 occurred a further 5.7 years later (mean age=67.6, SD=2.5, range=61.4 to 73.3). All twin pairs were concordant for US military service at some time between 1965 and 1975. However, nearly 80% reported no combat experience. The sample is 88.3% white, 5.3% African-American, 3.4% Hispanic, and 3.0% “other” participants. Based on data from the US National Center for Health Statistics, the sample is very similar to American men in their age range with respect to health, education, and lifestyle characteristics ^[14]^. Written informed consent was obtained from all participants. The University of California San Diego and Boston University ethics committees approved the study. Study protocols were identical at each site.

### Biomarker data

Primary analyses focused on plasma-derived Aβ40, Aβ42, Aβ42/40, NFL and t-tau (p-tau was not available). All Wave 3 samples were collected under fasting conditions. Subjects began fasting at 9:00 pm the night before testing. The following morning between 8:00 am and 8:15 am, blood samples were acquired, frozen and stored at −80°C. Commercial kits were used to perform the biomarker concentration analysis. The Simoa Human Neurology 3-plex A (N3PA) Immunoassay was used to measure Aβ40, Aβ42, and t-tau, while the Simoa NF-light assay was used to measure NFL. Standard exclusion criteria included hemolysis, subjects with mean NFL > 100, mean t-tau > 80, mean Aβ42/40 > 0.20, or a coefficient of variance > 20%. All biomarker assays were performed in Dr. Rissman’s laboratory at the University of California, San Diego.

Among subjects with complete biomarker data, 14.6%, 79.9%, and 5.5% were assessed in Boston, San Diego, and in their hometowns, respectively. A total of 80% of twin pairs were assessed on the same day. The average storage time between collection and processing was 1.9 years (SD=0.72).

The effects on each biomarker of age at assessment, testing site, storage time, ethnicity, and whether or not twins pairs were assessed on the same day were estimated and removed using the *umx_residualize*() function within the umx software package ^[65]^. All scores were then log-transformed in R_4.0.3 [61]_ to reduce skewness prior to our model fitting. Results from *umx_residualize*() revealed that prior to residualization, age at blood draw was not associated with Aβ40, Aβ42, and tau. It was however, linked to higher NFL (β = 26.12, t = 2.823, p = 0.005). In terms of location, San Diego subjects had significantly higher levels of Aβ40 (β = 88.45, t = 8.592, p < 0.001) and Aβ42 (β = 4.71, t = 10.897, p < 0.001). Longer storage time was significantly related to lower Aβ40 (β = −1560.39, t = −5.006, p < 0.001), and Aβ42 (β = −49.34, t = −3.808, p < 0.001) and higher t-tau levels (β = 18.40, t = 3.186, p = 0.001). Finally, neither self-reported ethnicity nor being concordant for assessment day (number of days measured apart) were associated with individual differences in any of the biomarkers.

### Statistical Analyses

The OpenMx_2.9.9.1_ software package ^[21]^ in R_3.4.1_^[61]^ was used to estimate twin pair correlations and to fit univariate and multivariate genetic twin models ^[7]^.

### Univariate analyses

In univariate twin analyses, the total variation in each biomarker was decomposed into additive genetic (A), shared or common environmental (C), and unshared or unique environmental (E) variance components (see Figure 1). This approach is referred to as the ‘ACE’ variance component model. The decomposition is achieved by exploiting the expected genetic and environmental correlations between monozygotic (MZ) and dizygotic (DZ) twin pairs. MZ twin pairs are genetically identical, whereas DZ twin pairs share, on average, half of their genes. Therefore, the MZ and DZ twin pair correlations for the additive genetic effects are fixed to r_A_=1.0 and r_A_=0.5 respectively. The modelling assumes that the sharing of environmental effects (C) is equal in MZ and DZ twin pairs (r_C_=1.0), while unshared environmental effects (E) are by definition uncorrelated and include measurement error.

**Figure 1.**
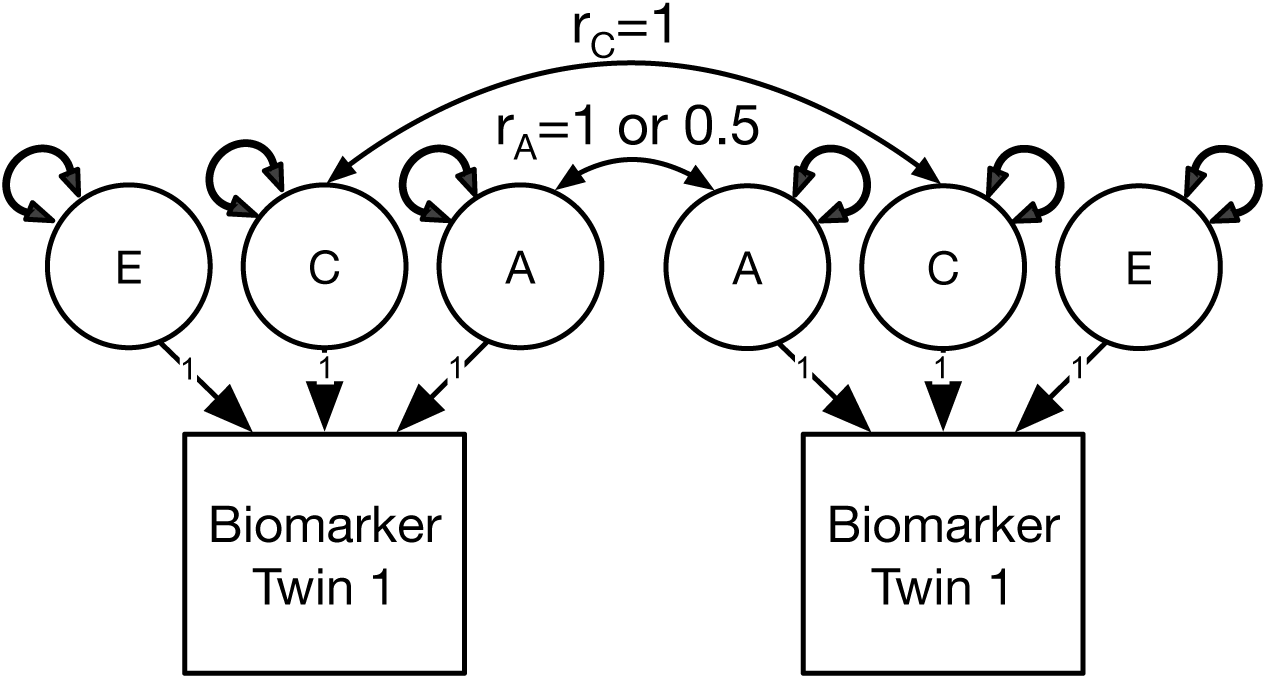
Univariate variance decomposition to estimate the relative contribution of genetic & environmental influences in each biomarker Note: A = additive genetic, C = common or shared environmental, & E = unshared environmental influences. r_C_ = correlation of 1 for MZ and DZ twin pairs. r_A_ = 1 or 0.5 for MZ & DZ twin pairs respectively.

### Multivariate analyses to test competing theories

This univariate method was extended to the multivariate case to estimate the significance of genetic and environmental influences within and shared between each of the biomarkers. To provide a reference for contrasting competing genetic and environmental models, we first fitted a multivariate ACE ‘correlated factors’ model ^[79]^ (Figure 2) before successively dropping the A and C components of variance to determine the best overall fit to the data.

**Figure 2.**
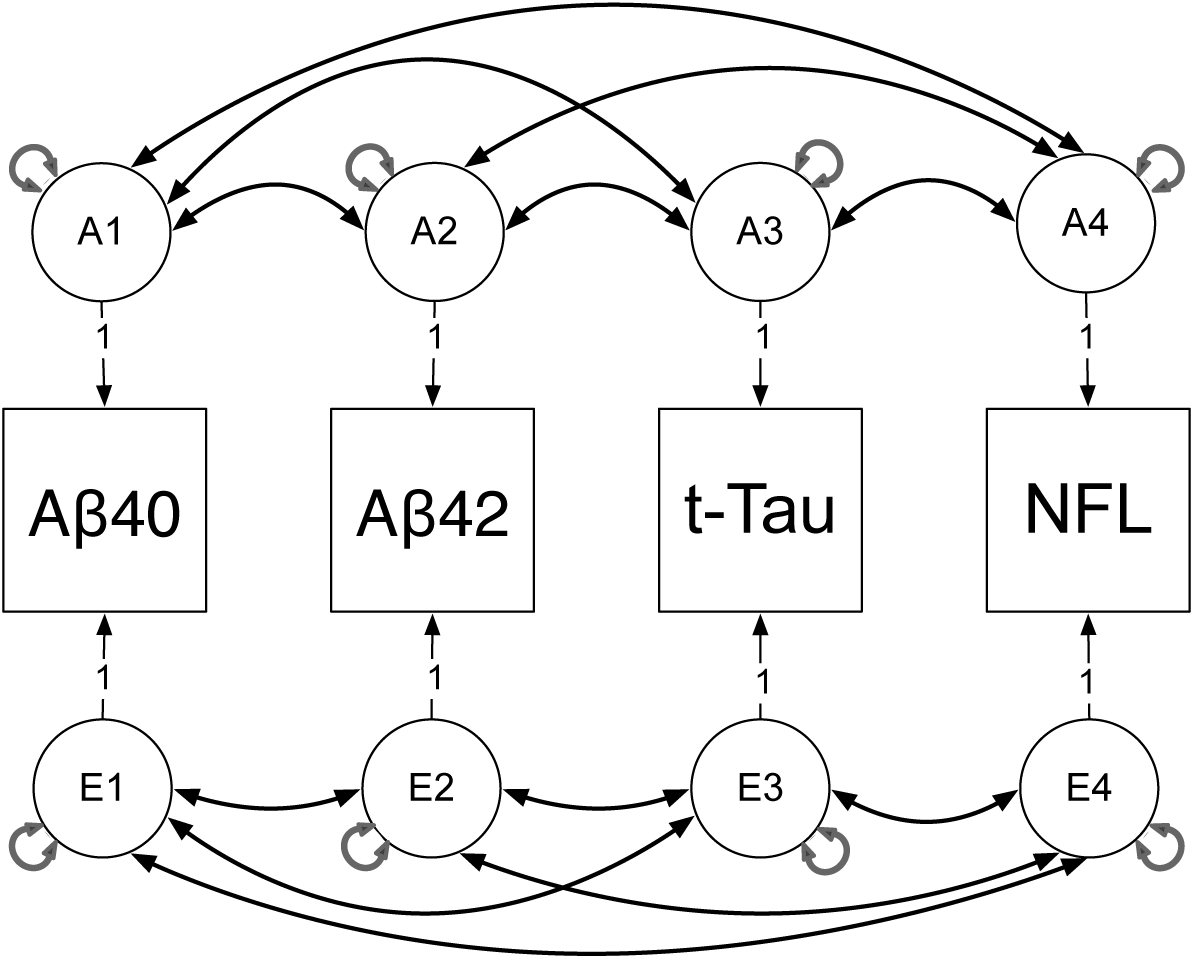
Multivariate correlated liabilities model to estimate the sources of genetic & environmental variances and covariances between the Aβ40, Aβ42, t-Tau & Neurofilament Light (NFL) biomarkers. Note: A1-A4 & E1-E4 denote latent additive genetic & non-shared environmental risk factors for the 5 biomarkers. Latent shared environmental factors not shown for brevity. Double-head arrows denote variances & covariances within & between latent factors.

### Model fit

For the univariate and multivariate analyses, we determined the most likely sources of variance by fitting three additional sub-models in which the i) C, ii) A, and iii) C and A influences were fixed to zero. In other words, we tested the statistical likelihood of the AE, CE, and E models, respectively. The significance of the A, C and E parameters was determined using the change in the minus two Log-Likelihood (ι1-2LL). Under certain regularity conditions, the ι1-2LL is asymptotically distributed as chi-squared with degrees of freedom equal to the difference in the number of free parameters in the two models. The determination of the best-fitting model was also based on the optimal balance of complexity and explanatory power by using Akaike’s Information Criterion (AIC) ^[2]^.

## Results

Table 1 shows the numbers of complete and incomplete twins by zygosity for each biomarker.

**Table 1.** Descriptive statistics including numbers of complete twin pairs & singletons for each of the five bio-markers by zygosity.

|  | N | Monozygotic |  | Dizygotic |  |
| --- | --- | --- | --- | --- | --- |
|  |  | Complete | Singletons | Complete | Singletons |
| 1. Aβ40 | 1015 | 240 | 109 | 159 | 108 |
| 2. Aβ42 | 998 | 237 | 104 | 157 | 106 |
| 3. Aβ42/40 | 998 | 237 | 104 | 157 | 106 |
| 4. t-tau | 963 | 213 | 132 | 142 | 121 |
| 5. NFL | 1052 | 257 | 100 | 171 | 96 |

### Testing the assumption of mean and variance homogeneity

Prior to the twin modelling of the combined MZ and DZ twin data we tested the assumption of mean and variance homogeneity for each biomarker using the residualized data. Supplementary Table S1 shows all mean and variance parameter estimates for the fully saturated and the constrained homogeneity models. As shown in Supplementary Table S2, constraining the means and variances to be equal within twin pairs and across zygosity resulted in a significant change in chi-square for Aβ40 and t-tau using a Bonferroni corrected p-value of p=0.01. Efforts to transform these two biomarkers or eliminate outliers using the Winsorize, Interquartile Range, and Box-Cox procedures did not alter this pattern of results. This was likely due to the small numbers of complete and incomplete twin pairs within each zygosity group. Notwithstanding this limitation, all subsequent analyses proceeded under the assumption of mean variance homogeneity for each biomarker.

### Strength of association

The phenotypic correlations and their 95% confidence intervals are shown in Table 2. Since Aβ42/40 is a linear function of Aβ42 and Aβ40, correlations between each of the Aβ biomarkers and the ratio were not calculated. Of note was the very high phenotypic correlation between Aβ40 and Aβ42. Both the Aβ42 and t-tau and the Aβ40 and t-tau correlations were not significant, and although the correlation between Aβ42/40 and t-tau was significant it was very small. The three Aβ biomarker correlations with NLF were significant but small. Likewise, the association between t-tau and NFL was significant but small.

**Table 2.** Pairwise polyserial phenotypic correlations & their standard errors between the five bio-markers along with the monozygotic (r_MZ_) & dizygotic (r_DZ_) twin pair correlations (including 95% confidence intervals).

|  | Phenotypic correlations (95%CI) |  |  |  |  | Twin pair correlations* |  |
| --- | --- | --- | --- | --- | --- | --- | --- |
| | 1 | 2 | 3 | 4 | 5 | $r_{MZ}$ (95%CI) | $r_{DZ}$ (95%CI) |
| 1. Aβ40 | 1 | | | | | 0.75 (0.68, 0.80) | 0.59 (0.43, 0.70) |
| 2. Aβ42 | 0.88 ( 0.86, 0.89) | 1 | | | | 0.49 (0.39, 0.58) | 0.27 (0.10, 0.42) |
| 3. Aβ42/40 | - | - | 1 | | | 0.39 (0.27, 0.49) | 0.26 (0.08, 0.41) |
| 4. t-tau | -0.08 (-0.14, -0.01) | 0.06 (-0.01, 0.12) | 0.08 (0.01, 0.10) | 1 |  | 0.60 (0.49, 0.67) | 0.24 (0.09, 0.37) |
| 5. NFL | 0.21 ( 0.15, 0.27) | 0.28 ( 0.22, 0.34) | 0.20 (0.16, 0.21) | 0.17 (0.11, 0.23) | 1 | 0.55 (0.46, 0.62) | 0.27 (0.11, 0.42) |

Note: \*Twin pair correlations calculated under the assumption of mean and variance homogeneity.

### Twin pair correlations

Table 2 also shows the twin pair correlations by zygosity for each biomarker. When familial aggregation is entirely attributable to shared family environments, the MZ and DZ twin pair correlations are expected to be statistically equivalent. In contrast, when familial aggregation is driven entirely by additive genetic factors, DZ twin pair correlations will be ½ the size (or less in the presence of genetic non-additivity) the MZ twin pair correlations.

For Aβ40, the DZ twin pair correlation was greater than ½ the MZ correlation suggesting a combination of additive genetic and shared environmental factors driving familial aggregation. For all remaining biomarkers, the DZ twin pair correlations were approximately ½ their MZ twin pair counterparts. This pattern of correlations is consistent with the hypothesis that additive genetic factors alone explain familial aggregation, while all remaining variance is attributable to aspects of the environment unshared between siblings.

### Univariate analyses

Table 3 summarizes the best fitting univariate models. Detailed model fitting results are shown in Supplementary Table S3.

**Table 3.**
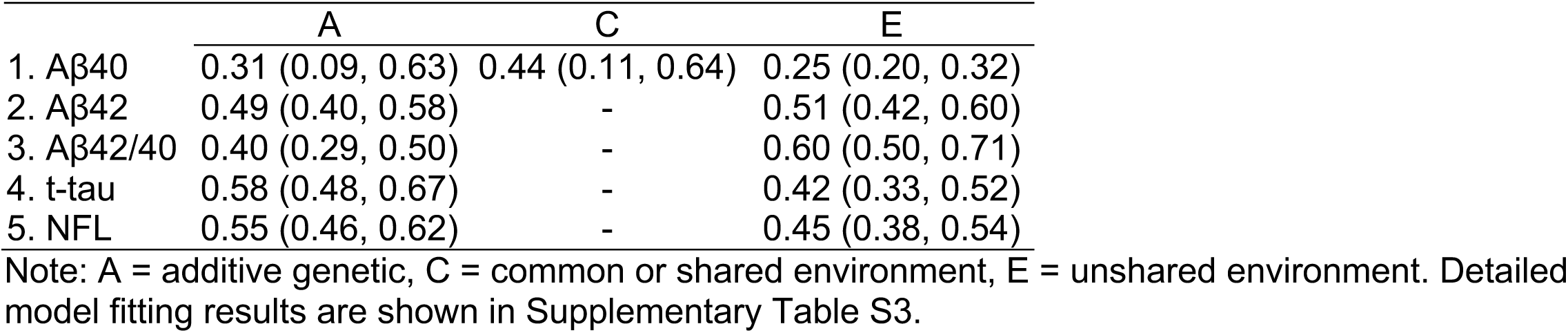
Standardized estimates of the additive genetic (A), shared (C) & unshared (E) environmental influences (including 95% confidence intervals) under the best fitting univariate models.

### Aβ40

Based on the lowest AIC and the significant changes in the Δ-2LL associated with the AE, CE and E sub-models, the full ACE model was identified as the best fitting. Consistent with the pattern of MZ and DZ twin pair correlations, familial aggregation was associated with a combination of additive genetic (31%) and shared environmental (44%) influences accounting for three quarters of the total variance in this biomarker.

### Aβ42

Based on the lowest Akaike’s Information Criterion (AIC) and a non-significant change in Δ-2LL, the AE sub-model was chosen as the best fitting. Here, familial aggregation could be entirely explained by additive genetics alone, which in turn accounted 49% of all individual differences in this biomarker.

### Aβ42/40

Similar to Aβ42, the AE sub-model again provided the best fit to the data with additive genetic variance accounting for 40% of the total variance in this biomarker.

### t-tau

The AE sub-model did not deteriorate significantly when all shared environmental effects (C) were removed. This model also yielded the lowest AIC value. Here, familial aggregation was entirely explained by additive genetics alone, which accounted for 58% of the total variance in this biomarker.

### NFL

The AE sub-model again provided the best fit to the data, with additive genetic influences accounting for all familial aggregation, which in turn explained 55% of all individual differences in this biomarker.

### Multivariate analyses

Multivariate analyses were next used to estimate the size and significance of the genetic and environmental influences within and between the four biomarkers (Aβ42, Aβ40, t-tau and NFL). The Aβ42/40 biomarker was not modelled since this ratio measure is a linear combination of Aβ42 and Aβ40. As shown in Table 4, the AE, CE and E sub-models all deteriorated significantly when compared to the full ACE model.

**Table 4.**
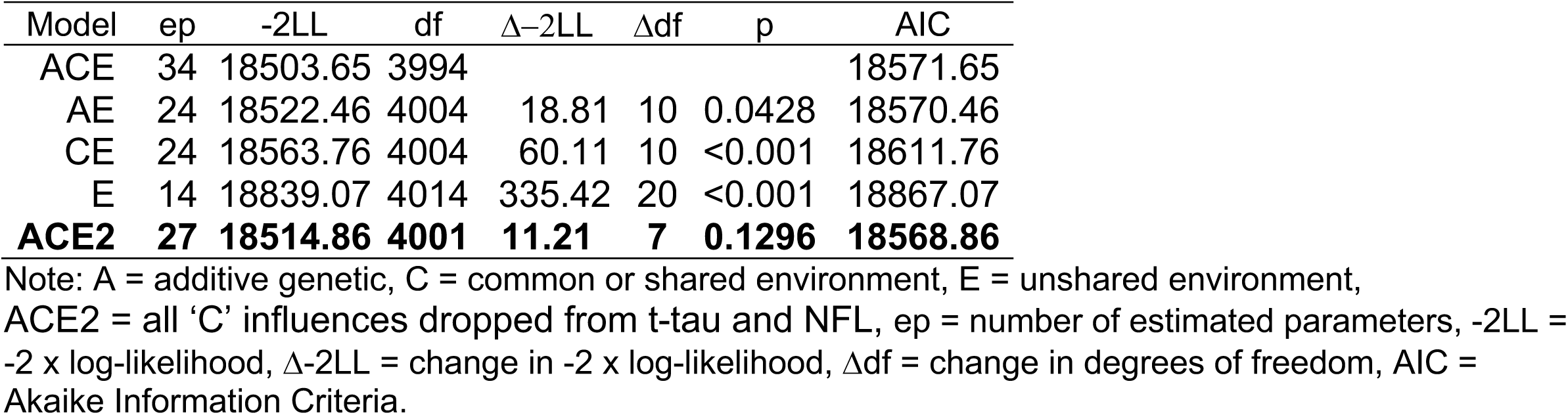
Multivariate model fitting comparisons between the reference ACE and the AE, CE, E and ACE2 nested sub-models. The best fitting model in bold font.

Under the best fitting ACE model, the shared environmental factor correlations (r_c_) between either t-tau or NFL and the two Aβ biomarkers were undefined i.e., empirically under-identified. This can arise when ‘C’ influences on one or more traits are estimated to be zero. Therefore, based on the twin pair correlations and the univariate results, we removed all ‘C’ influences from t-tau and NFL before re-running. This model, labeled ‘ACE2’ in Tables 4–5, provided the best overall fit to the data as judged by the non-significant change in chi-square and marginally lowest AIC.

**Table 5.**
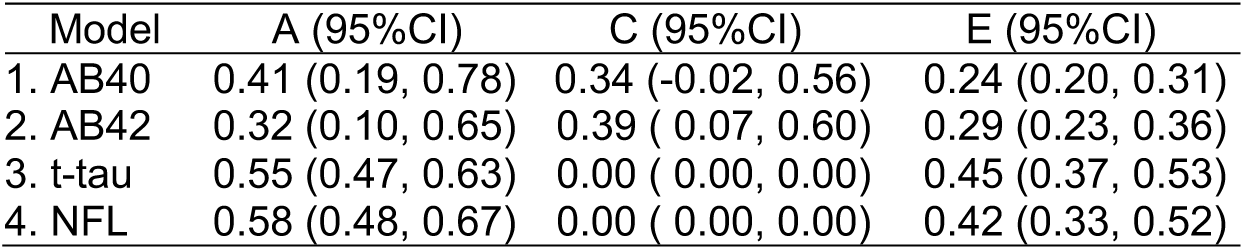
Standardized estimates of the additive genetic & unshared environmental influences for the multivariate ACE and better fitting ACE2 model.

Under ‘ACE2’, familial aggregation in Aβ40 was explained by a combination of additive genetic (41%) and shared environmental (34%) influences (see Table 5). We note, however, that these ‘C’ influences were non-significant. Next, familial aggregation in Aβ42 was explained by a combination of significant additive genetic (32%) and shared environmental (39%) influences. For t-tau and NFL, familial aggregation was entirely explained by additive genetic influences ranging 55% to 58%. For all four biomarkers, all remaining sources of variation were explained by unshared environmental influences including measurement error.

Table 6 summarizes the multivariate genetic and environmental correlations between the four biomarkers based on the ACE2 best fitting model. Genetically, Aβ40 and Aβ42 were statistically identical (r = 1.00). All other genetic factor correlations were significant but small: Aβ40 correlated negatively with t-tau (r_a_ = −0.19); Aβ42 correlated positively with NFL (r_a_ = 0.21); while t-tau and NFL correlated positively (r_a_ = 0.21). The shared environmental factor correlation between Aβ40 and Aβ42 was non-significant. In terms of unshared environmental correlations, there was a moderate correlation between Aβ40 and Aβ42 (r_e_ = 0.66). Neither of the Aβ biomarker shared any significant ‘E’ influences with t-tau. In contrast, there were significant but small positive environmental correlations between each of the Aβ biomarkers and NFL ranging from r_e_ = 0.28 to r_e_ = 0.38. Finally, there was a significant but small unshared environmental correlation between t-tau and NFL (r_e_ = 0.18).

**Table 6.** Additive genetic, shared & unshared (or unique) environmental latent factor correlations based on the best fitting multivariate ‘ACE2’ model.

| Genetic correlations |  |  |  |  |
| --- | --- | --- | --- | --- |
|  | Aβ40 | Aβ42 | t-tau | NFL |
| Aβ40 | 1.00 |  |  |  |
| Aβ42 | 1.00 ( 0.92, 1.00) | 1.00 |  |  |
| t-tau | 0.12 (-0.05, 0.30) | 0.21 ( 0.02, 0.45) | 1.00 |  |
| NFL | -0.19 (-0.41,-0.02) | -0.12 (-0.38, 0.09) | 0.18 ( 0.03, 0.32) | 1.00 |
| Shared environmental correlations |  |  |  |  |
|  | Aβ40 | Aβ42 | t-tau | NFL |
| Aβ40 | 1.00 |  |  |  |
| Aβ42 | 0.91 (-1.00, 1.00) | 1.00 |  |  |
| t-tau | - | - | 1.00 |  |
| NFL | - | - | - | 1.00 |
| Unshared environmental correlations |  |  |  |  |
|  | Aβ40 | Aβ42 | t-tau | NFL |
| Aβ40 | 1.00 |  |  |  |
| Aβ42 | 0.66 ( 0.59, 0.73) | 1.00 |  |  |
| t-tau | 0.05 (-0.09, 0.19) | 0.13 ( 0.00, 0.27) | 1.00 |  |
| NFL | 0.38 ( 0.26, 0.48) | 0.28 ( 0.17, 0.39) | 0.18 ( 0.04, 0.30) | 1.00 |
| Additive genetic correlations |  |  |  |  |
|  | Aβ4240 | t-tau | NFL |  |
| Aβ4240 | 1 |  |  |  |
| t-tau | 0.05 (-0.14,0.06) | 1 |  |  |
| NFL | 0.29 ( 0.12,0.44) | 0.18 ( 0.03,0.32) | 1 |  |
| Unshared environmental correlations |  |  |  |  |
|  | Aβ4240 | t-tau | NFL |  |
| Aβ42 | 1 |  |  |  |
| t-tau | 0.11 ( 0.07,0.12) | 1 |  |  |
| NFL | 0.14 ( 0.13,0.25) | 0.17 ( 0.04,0.30) | 1 |  |

## Discussion

To our knowledge, this is the first study examining the genetic and environmental etiology of AD-related biomarkers in blood plasma. We estimated the relative contribution of genetic and environmental influences on five biomarkers using biometrical genetic twin models. Our univariate modelling revealed significant familial aggregation attributable to additive genetic and shared environmental influences. Additive genetics explained 31% to 58% of the total variances in Aβ40, Aβ42, Aβ42/40, t-tau and NFL. The univariate analyses also revealed that shared environmental influences explained 44% of the total variance in Aβ40. We then employed the statistically more powerful multivariate twin analyses to explore the role of genetic and environmental influences within and between the biomarkers. Shared environmental effects explained 34% to 39% of the total variance in Aβ biomarkers, but were non-significant in Aβ40. Additive genetics explained 32% to 41% of the variance in Aβ42 and Aβ40 respectively. In contrast, familial aggregation in t-tau and NFL was entirely explained by additive genetic influences, which accounted for over one-half of the total variance ranging 55% to 58%. Thus, individual differences in AD-related plasma biomarkers are substantial and can be explained by varying combinations of genetic and shared environmental influences.

The multivariate analyses revealed noteworthy genetic and environmental correlations between the plasma biomarkers. For instance, the genetic correlation between Aβ40 and Aβ42 biomarkers, which are the two most predominant species of Aβ in humans ^[33]^, suggest that their genetic influences are identical. Indeed, the very high genetic correlation is commensurate with the fact that Aβ42 is only two amino acids longer than Aβ40, and can be distinguished by the presence of an Ile–Ala dipeptide at the C-terminal end of an otherwise identical 40 amino acid peptide ^[1, 33]^. Despite our results, Aβ42 has been shown to be more strongly linked to AD pathology than is Aβ40^[8–10]^. In vitro findings have shown how Aβ42 has higher fibril nucleation and elongation rates, as well as larger oligomers and more toxic assemblies than Aβ40 ^[33]^. We speculate that a lower genetic correlation might be observed in CSF, or that independent genetic influences might be detectable in neuronally derived Aβ40 and Aβ42, or at later stages of AD progression.

According to the amyloid cascade hypothesis ^[5, 6]^, Aβ aggregation drives the accumulation of tau tangles, resulting in synaptic dysfunction, neurodegeneration and progression to cognitive decline. If Aβ aggregation causally impacts t-tau via genetic mechanisms, then significant genetic covariance between the Aβ and t-tau biomarkers should be observed. Our results lend partial support to this model; the genetic correlation (r_g_) between t-tau and Aβ42 was significant, and although the Aβ40 and t-tau r_g_ was non-significant, the upper bound 95% confidence interval was +0.30. Given the Aβ4240 is a linear function of Aβ40 and Aβ42, we ran a *post-hoc* tri-variate analysis of Aβ4240, t-tau and NFL to validate this trend. Here, the genetic correlation between Aβ4240 and t-tau was small and non-significant with a low upper bound (r_g_ = 0.05 [95%CI = −0.14,0.06]). In contrast, the r_g_ between Aβ4240 was NFL was larger and significant (r_g_ = 0.29 [95%CI = 0.12,0.44]). Also noteworthy was that NFL having the highest heritability point estimate (58%). NFL is a marker of axonal damage ^[11]^ and plasma NFL has been significantly linked to neurodegeneration ^[70, 85]^. Our ongoing fourth wave follow-up assessment (which is collecting both t- and p-tau) will reveal if the genetic correlations between the Aβs and tau (p- and t-tau) or the Aβs and NFL increase and reach significance when the mean age of the sample is projected to be 74 years.

In terms of random environmental influences unshared between siblings, these effects explained less than one-half of the standardized variance in each biomarker. As mentioned previously, ‘E’ effects necessarily include measurement error. Therefore, the multivariate unshared environmental correlations will also capture correlated measurement errors. If the assay contributed to similar measurement errors across biomarkers, then provided the measurement errors were uncorrelated between siblings, this would have resulted in more uniform unshared environmental correlations. Despite Aβ40, Aβ42 and t-tau each being measured on the same N3PA assay, the pattern of unshared environmental correlations was not uniform. For instance, there was a moderate significant r_e_ between Aβ40 and Aβ42 versus small non-significant correlations between the Aβ biomarkers and t-tau. Regarding the unshared environmental correlations between NFL and the three other biomarkers, we caution against over-interpreting the significance of the small unshared environmental correlations since the overall magnitude of their phenotypic associations was small (see Table 2). It is possible that the observed pattern of unshared environmental correlations arose, in part, from differential rates of dilution, degradation, or metabolism effecting the biomarkers after entering the bloodstream, which could have introduced additional variance unshared between siblings.

We also note the discrepancy between the univariate and multivariate variance components for the Aβ biomarkers. When estimating univariate variance components, information is derived solely from the within-trait cross-twin covariance structure. In contrast, the multivariate variance component estimates rely upon additional information from the cross-twin cross-trait covariances. Inspection of the multivariate phenotypic cross-twin cross-trait correlations (See Supplementary Table S4) reveals that both DZ Aβ cross-twin cross-trait correlations were greater than ½ their MZ Aβ cross-twin cross-trait correlations. This is consistent with the shared environmental influences on the Aβ biomarkers observed in the multivariate results. Given the increased power of multivariate twin models and the use of the correlated factors model to obtain the correct Type 1 error rate ^[79]^, the univariate Aβ42 results (Table 2) may represent outliers when compared to the multivariate data.

Broadly, our findings have implications in terms of the ATN framework

^[42, 57]^ that relies on imaging and CSF biomarkers divided into three binary classes: Aβ biomarkers (A); tau pathology (T); biomarkers measuring neurodegeneration or neuronal injury (N). Notwithstanding the limitations of this approach ^[55, 56, 78]^, this framework was intended to be flexible in terms of adding either new biomarkers or entire classes of new biomarkers ^[57]^. Our demonstration of significant heritability and genetic covariance suggests that that the inclusion of plasma biomarkers within the ATN framework may be warranted. Indeed, Koycheva’s meta-analysis of 83 phenotypic studies highlighted the validity of ATN plasma biomarkers (when measured using ultrasensitive techniques) to differentiate significantly between AD patients and controls ^[83]^. Of course, practical implementation would require agreed upon positivity cutoffs for plasma biomarkers. Our next step will be to determine the degree to which the genetic and environmental variances in our plasma biomarkers can reliably predict individual differences in MCI and risk of AD.

It is important to note that the correlations between Aβ and t-tau were positive. This may seem rather counterintuitive given that, like CSF Aβ, lower plasma Aβ levels are generally considered to be more pathological ^[52][68]^. However, pattern is consistent with results based on at least 4 independent samples that have revealed a quadratic association between CSF Aβ in cognitively normal adults ^[52][68]^, whereby the association is younger or cognitively normal individuals (inverted-U pattern suggests an early increase in CSF Aβ production followed later by a sequestration in amyloid plaques, while CSF tau increases throughout). In our largely cognitively normal sample with a mean age of only 68, most individuals may be on the rising/lower side of the inverted-U curve for plasma Aβ. As such, we might expect the correlations between Aβ and tau to switch from positive to negative in the next wave of the study.

## Limitations

Our results should be interpreted in the context of potential limitations.

First, we explored only a limited number of plasma biomarkers on existing arrays. Although we plan to obtain them from remaining samples, we did not have measures of p-tau. Three p-tau isoforms (181, 217, and 231) have, for example, been shown to predict amyloidosis and progression to AD ^[82]^. The genetic etiology of these isoforms remains undetermined including their covariance with the Aβ and NFL biomarkers. As noted, t-tau is not generally considered as good an indicator of neurofibrillary tangles as p-tau, so it is probably not the ideal marker of T in the ATN framework. On the other hand, t-tau and p-tau are very highly correlated.

Second, because between-group differences are likely to be complex ^[86]^, our results may not generalize to women or to other ancestral groups. Due in part to their greater longevity, women are disproportionately affected by AD in terms of both disease prevalence and severity ^[30]^.

Therefore, it is unclear to what extent there are sex differences in the means and variance components of these plasma-based biomarkers in similarly aged women, and whether such differences translate into different outcomes. Regarding ancestral differences, African Americans are at greater risk of developing AD compared to Caucasians ^[64, 84]^. In terms of specific biomarkers, African-Americans have lower levels of CSF tau that appear to be unrelated to neurodegeneration ^[47, 67, 71]^, and when compared to Caucasians, African-Americans with the *APOE*-ε4 risk allele also have lower CSF t-tau and p-tau181 ^[71]^. Therefore, it is plausible that the genetic and environmental etiologies of the plasma biomarkers differ between ancestral groups. Only by ascertaining larger and ancestrally varied samples can we begin to test hypotheses regarding important group differences, including the generalizability and validity of the overall ATN framework.

These above limitations are offset by notable strengths. Among subjects with biomarker data, the mean level of education was 13.99 years (SD=2.08), which is similar to the general population for this age cohort. This is particularly important because low education is a known risk factor for AD ^[20]^. Additionally, some large biomarker studies have exclusion criteria for several health conditions, whereas the VETSA is a community-dwelling sample that does not exclude for these reasons. Therefore, the sample may also be more representative of at least men in their age group with respect to health factors.

### Conclusion

To our knowledge, this is the first study to explore the genetic and environmental influences in plasma AD-related biomarkers. In community-dwelling men at average age 68 years, these biomarkers are heritable. Genetic influences were associated with 32% to 41% of the variance in the Aβ biomarkers and over one-half of the variance in t-tau and NFL. The presence of ‘C’ in Aβ40 or Aβ42 implies that the impact of being reared together may be persistent in terms of influencing biomarker levels in early old age. Although the biomarkers examined here were not brain-derived, changes in plasma biomarkers occur at much the same time as their CSF counterparts ^[73]^, and are proving to be useful for screening individuals in the prodromal stages of AD ^[43, 74, 75]^. Future analyses should explore the sources of genetic and environmental covariance between plasma biomarkers, MCI, and risk of AD.

## Acknowledgements

This work was supported by the National Institute on Aging at the National Institutes of Health grant numbers R01s AG050595, AG022381, AG076838, AG037985; R25 AG043364, F31 AG064834; P01 AG055367, AG062483; and K01 AG063805. The funding sources had no role in the preparation, review, or approval of the manuscript, or the decision to submit the manuscript for publication. The U.S. Department of Veterans Affairs, Department of Defense; National Personnel Records Center, National Archives and Records Administration; Internal Revenue Service; National Opinion Research Center; National Research Council, National Academy of Sciences; and the Institute for Survey Research, Temple University provided invaluable assistance in the conduct of the VET Registry. The Cooperative Studies Program of the U.S. Department of Veterans Affairs provided financial support for development and maintenance of the Vietnam Era Twin Registry. We would also like to acknowledge the continued cooperation and participation of the members of the VET Registry and their families.

---

**Supplementary Table S1.** Monozygotic (MZ) and dizygotic (DZ) sample sizes, means, variances and covariances for i) the fully saturated model with 10 parameters and ii) the restricted mean and variance homogeneity model with 4 parameters.

| Fully saturated model<br>10 parameters (4 means + 4 variances + 2 covariances) per biomarker |  |  |  |  | Mean & variance homogeneity model<br>4 parameters (1 mean + 1 variance + 2 covariances) per biomarker |  |  |
| --- | --- | --- | --- | --- | --- | --- | --- |
|  | N | Means (SE) | Variances & covariances |  | Means (SE) | Variances & covariances |  |
|  |  |  | Twin1 | Twin2 |  | Twin1 | Twin2 |
| 1. Aβ40 |  |  |  |  |  |  |  |
| MZ Twin 1 | 294 | 54.22 (0.29) | 27.63 (23.51 - 32.77) |  | 54.61 (0.16) | 17.96 (16.15 - 20.06) |  |
| MZ Twin 2 | 295 | 54.66 (0.26) | 19.87 (16.35 - 24.04) | 21.68 (17.92 - 26.42) |  | 13.39 (11.31 - 15.68) | 17.96 (16.15 - 20.06) |
| DZ Twin 1 | 216 | 54.75 (0.22) | 10.12 (8.38 - 12.41) |  |  | 17.96 (16.15 - 20.06) |  |
| DZ Twin 2 | 210 | 54.85 (0.20) | 2.37 (0.37 - 4.40) | 8.85 ( 7.35 - 10.78) |  | 10.62 ( 7.27 - 13.46) | 17.96 (16.15 - 20.06) |
| 2. Aβ42 |  |  |  |  |  |  |  |
| MZ Twin 1 | 286 | 22.46 (0.19) | 10.40 (8.86 - 12.30) |  | 22.59 (0.11) | 8.86 (8.08 - 9.75) |  |
| MZ Twin 2 | 292 | 22.77 (0.17) | 5.06 (3.74 - 6.56) | 9.11 (7.74 - 10.84) |  | 4.35 (3.31 - 5.42) | 8.86 (8.08 - 9.75) |
| DZ Twin 1 | 213 | 22.53 (0.19) | 7.98 (6.62 - 9.73) |  |  | 8.86 (8.08 - 9.75) |  |
| DZ Twin 2 | 207 | 22.58 (0.19) | 1.85 (0.63 - 3.17) | 7.46 (6.20 - 9.10) |  | 2.42 (0.85 - 3.87) | 8.86 (8.08 - 9.75) |
| 3. Aβ42/40 |  |  |  |  |  |  |  |
| MZ Twin 1 | 286 | 20.41 (0.00) | 0.0020 (0.0017 - 0.0024) |  | 20.41 (0.00) | 0.0017 (0.0015 - 0.0018) |  |
| MZ Twin 2 | 292 | 20.41 (0.00) | 0.0008 (0.0005 - 0.0011) | 0.0017 (0.0036 - 0.0021) |  | 0.0007 (0.0004 - 0.0009) | 0.0017 (0.0015 - 0.0018) |
| DZ Twin 1 | 213 | 20.41 (0.00) | 0.0014 (0.0012 - 0.0017) |  |  | 0.0017 (0.0015 - 0.0018) |  |
| DZ Twin 2 | 207 | 20.41 (0.00) | 0.0003 (0.0001 - 0.0005) | 0.0014 (0.0014 - 0.0038) |  | 0.0004 (0.0001 - 0.0007) | 0.0017 (0.0015 - 0.0018) |
| 4. t-tau |  |  |  |  |  |  |  |
| MZ Twin 1 | 271 | 22.90 (0.05) | 0.81 (0.69 - 0.97) |  | 22.97 (0.04) | 1.21 (1.10 - 1.34) |  |
| MZ Twin 2 | 287 | 22.93 (0.06) | 0.49 (0.36 - 0.64) | 1.10 (0.94 - 1.31) |  | 0.72 (0.57 - 0.87) | 1.21 (1.10 - 1.34) |
| DZ Twin 1 | 205 | 23.00 (0.09) | 1.56 (1.29 - 1.90) |  |  | 1.21 (1.10 - 1.34) |  |
| DZ Twin 2 | 200 | 23.07 (0.08) | 0.42 (0.16 - 0.70) | 1.41 (1.16 - 1.73) |  | 0.29 (0.11 - 0.46) | 1.21 (1.10 - 1.34) |
| 5. NFL |  |  |  |  |  |  |  |
| MZ Twin 1 | 303 | 23.52 (0.31) | 29.91 (25.57 - 35.30) |  | 23.46 (0.18) | 25.73 (23.50 - 28.28) |  |
| MZ Twin 2 | 311 | 23.73 (0.28) | 15.57 (12.16 - 19.53) | 24.71 (21.19 - 29.04) |  | 14.15 (11.40 - 17.05) | 25.73 (23.50 - 28.28) |
| DZ Twin 1 | 219 | 23.44 (0.36) | 28.22 (23.50 - 34.29) |  |  | 25.73 (23.50 - 28.28) |  |
| DZ Twin 2 | 219 | 23.13 (0.30) | 6.17 ( 2.41 - 10.21) | 19.51 (16.26 - 23.70) |  | 7.07 ( 2.72 - 11.13) | 25.73 (23.50 - 28.28) |

**Supplementary Table S2.** Change in model fit associated with the comparison between a fully saturated model with 10 parameters per biomarker (4 means, 4 variances and 2 covariances) and a ‘mean and variance homogeneity’ model with just 4 parameters (1 mean, 1 variance and 2 covariances). Also shown are the Comparative fit index (CFI), Tucker Lewis index (TLI), and Root Mean Square Error of Approximation (RMSEA) statistics for the constrained 4-parameter ‘mean and variance homogeneity’ model. The CFI, TLI and RMSEA were derived using the mxRefModels option in OpenMx.

| | Δ-2LL | Δdf | p | CFI | TLI | RMSEA |
| --- | --- | --- | --- | --- | --- | --- |
| 1. Aβ40 | 95.84 | 6 | <0.001 | 0.23 | 0.74 | 0.13 |
| 2. Aβ42 | 10.73 | 6 | 0.097 | 0.93 | 0.98 | 0.03 |
| 3. Aβ42/40 | 11.84 | 6 | 0.066 | 0.85 | 0.95 | 0.03 |
| 4. t-tau | 30.47 | 6 | <0.001 | 0.63 | 0.88 | 0.07 |
| 5. NFL | 15.52 | 6 | 0.017 | 0.90 | 0.97 | 0.04 |

Note: Δ-2LL = change in -2 x log-likelihood, $\Delta$ df = change in degrees of freedom.

**Supplementary Table S3.** Univariate model fitting comparisons & the standardized estimates of the additive genetic (A), shared (C) & unshared (E) environmental influences (including 95% confidence intervals) under the competing ACE, AE, CE & E models. Best fitting model in bold font.

| | Model | ep | -2LL | df | Δ-2LL | Δdf | p | AIC | A | C | E |
| --- | --- | --- | --- | --- | --- | --- | --- | --- | --- | --- | --- |
| 1. Aβ40 | **ACE** | **4** | **5548.60** | **1011** | | | | **5556.60** | **0.31 (0.09, 0.63)** | **0.44 (0.11, 0.64)** | **0.25 (0.20, 0.32)** |
|  | AE | 3 | 5554.58 | 1012 | 5.98 | 1 | 0.0145 | 5560.58 | 0.74 (0.67, 0.79) | - | 0.26 (0.21, 0.33) |
|  | CE | 3 | 5556.47 | 1012 | 7.88 | 1 | 0.0050 | 5562.47 | - | 0.69 (0.62, 0.75) | 0.31 (0.25, 0.38) |
|  | E | 2 | 5659.29 | 1013 | 110.70 | 2 | 0.0000 | 5663.29 | - | - | 1.00 |
| 2. Aβ42 | ACE | 4 | 4931.92 | 994 | | | | 4939.92 | 0.43 (0.08, 0.82) | 0.06 (-0.31, 0.36) | 0.51 (0.43, 0.61) |
|  | **AE** | **3** | **4932.03** | **995** | **0.11** | **1** | **0.7450** | **4938.03** | **0.49 (0.40, 0.58)** | - | **0.51 (0.42, 0.60)** |
|  | CE | 3 | 4937.98 | 995 | 6.06 | 1 | 0.0139 | 4943.98 | - | 0.41 (0.32, 0.50) | 0.59 (0.50, 0.68) |
|  | E | 2 | 4999.80 | 996 | 67.88 | 2 | 0.0000 | 5003.80 | - | - | 1.00 |
| 3. Aβ42/40 | ACE | 4 | -3600.06 | 994 | | | | -3592.06 | 0.26 (-0.11, 0.67) | 0.13 (-0.24, 0.44) | 0.61 (0.51, 0.73) |
|  | **AE** | **3** | **-3599.56** | **995** | **0.50** | **1** | **0.4780** | **-3593.56** | **0.40 (0.29, 0.50)** | - | **0.60 (0.50, 0.71)** |
|  | CE | 3 | -3544.23 | 995 | 55.83 | 1 | 0.0000 | -3538.23 | - | 0.17 (-0.08, 0.64) | 0.83 (0.36, 1.08) |
|  | E | 2 | -3418.32 | 996 | 181.74 | 2 | 0.0000 | -3414.32 | - | - | 1.00 |
| 4. t-tau | ACE | 4 | 2816.08 | 959 |  |  |  | 2824.08 | 0.72 (0.39, 1.05) | -0.12 (-0.42, 0.16) | 0.40 (0.33, 0.51) |
|  | **AE** | **3** | **2816.76** | **960** | **0.68** | **1** | **0.4091** | **2822.76** | **0.58 (0.48, 0.67)** | - | **0.42 (0.33, 0.52)** |
|  | CE | 3 | 2834.65 | 960 | 18.57 | 1 | 0.0000 | 2840.65 | - | 0.41 (0.31, 0.49) | 0.59 (0.51, 0.69) |
|  | E | 2 | 2887.80 | 961 | 71.72 | 2 | 0.0000 | 2891.80 | - | - | 1.00 |
| 5. NFL | ACE | 4 | 6295.86 | 1048 |  |  |  | 6303.86 | 0.55 (0.23, 0.91) | 0.00 (-0.34, 0.29) | 0.45 (0.38, 0.53) |
|  | **AE** | **3** | **6295.86** | **1049** | **0.00** | **1** | **0.9969** | **6301.86** | **0.55 (0.46, 0.62)** | - | **0.45 (0.38, 0.54)** |
|  | CE | 3 | 6307.87 | 1049 | 12.01 | 1 | 0.0005 | 6313.87 | - | 0.46 (0.37, 0.53) | 0.54 (0.47, 0.63) |
|  | E | 2 | 6395.86 | 1050 | 100.00 | 2 | 0.0000 | 6399.86 | - | - | 1.00 |

Note: ep = number of estimated parameters, -2LL = -2 x log-likelihood, Δ-2LL = change in -2 x log-likelihood, $\Delta$ df = change in degrees of freedom, AIC = Akaike Information Criteria.

**Supplementary Table S4.** Full Information Maximum Likelihood multivariate phenotypic polyserial correlations based on the full multivariate ACE model. Phenotypic MZ and DZ twin pair correlations are below and above the diagonal respectively. MZ and DZ cross-twin cross-trait correlations are shaded. All correlations were calculated under the assumption of mean and variance homogeneity within twin-pairs within variable.

|  | 1. | 2. | 3. | 4. | 5. | 6. | 7. | 8. |
| --- | --- | --- | --- | --- | --- | --- | --- | --- |
| 1. Twin 1 Aβ40 | 1.00 | 0.89 | 0.19 | -0.10 | 0.58 | 0.54 | 0.09 | -0.12 |
| 2. Twin 1 Aβ42 | 0.89 | 1.00 | 0.20 | -0.02 | 0.54 | 0.56 | 0.07 | -0.14 |
| 3. Twin 1 t-tau | 0.19 | 0.20 | 1.00 | 0.18 | 0.09 | 0.07 | 0.28 | 0.10 |
| 4. Twin 1 NFL | -0.10 | -0.02 | 0.18 | 1.00 | -0.12 | -0.14 | 0.10 | 0.25 |
| 5. Twin 2 Aβ40 | 0.75 | 0.72 | 0.06 | -0.11 | 1.00 | 0.89 | 0.19 | -0.10 |
| 6. Twin 2 Aβ42 | 0.72 | 0.72 | 0.10 | -0.05 | 0.89 | 1.00 | 0.20 | -0.02 |
| 7. Twin 2 t-tau | 0.06 | 0.10 | 0.55 | 0.10 | 0.19 | 0.20 | 1.00 | 0.18 |
| 8. Twin 2 NFL | -0.11 | -0.05 | 0.10 | 0.60 | -0.10 | -0.02 | 0.18 | 1.00 |

